# Reusable Microfluidic Chambers for Single-Molecule Microscopy

**DOI:** 10.1101/2024.09.27.615484

**Authors:** Janan Alfehaid, Sineth G. Kodikara, Tuqa Alhajri, Mohammad Lutful Kabir, Hamza Balci

## Abstract

Maintaining a consistent environment in single molecule microfluidic chambers containing surface-bound molecules requires laborious cleaning and surface passivation procedures. Despite such efforts, variations in non-specific binding and background signals commonly occur across different chambers. Being able to reuse the chambers without degrading the surface promises significant practical and fundamental advantages; however, this necessitates removing the molecules attached to the surface, such as DNA, proteins, lipids or nanoparticles. Biotin-streptavidin attachment is widely used for such attachments as biotin can be readily incorporated to these molecules. In this study, we present single-molecule fluorescence experiments that demonstrate effective resetting and recycling of the chambers at least 10 times using photocleavable biotin (PC-biotin) and UV light exposure. This method differs from alternatives as it does not utilize any harsh chemical treatment of the surface. We show that all bound molecules (utilizing various PC-biotin attachment chemistries) can be removed from the surface by a 5-min UV exposure of a specific wavelength. Non-optimal wavelengths and light sources showed varying degrees of effectiveness. Our approach does not result in any detectable degradation of surface quality, as assessed by non-specific binding of fluorescently labeled DNA and protein samples, and recovery of DNA secondary structure and protein activity. The speed and efficiency of the resetting process, the cost-effectiveness of the procedure, and the widespread use of biotin-streptavidin attachment make this approach adaptable for a wide range of single-molecule applications.

## 1. Introduction

Reliable and reproducible surface-based single molecule measurements require indispensable surface characteristics, including negligible non-specific binding, low background signal, and minimal interaction with surface-attached molecules. To attain such surfaces, elaborate surface passivation protocols that employ bovine serum albumin (BSA), polyethylene glycol (PEG), or casein have been developed ^1–5^. In the most demanding applications, additional steps have been employed to further improve the surface quality. For example, adding a second coating of PEG with a significantly smaller molecular weight than the original coating (e.g., 5000 Da as the original and 333 Da as the second coating) increased the density of the polymer brush and reduced non-specific binding by ∼3-fold ^1^. Similarly, treating the PEG or BSA coated surfaces with a nonionic detergent (such as polysiloxane-Tween-20 or dimethyldichlorosilane-Tween-20) increases hydrophobicity of the surface and reduces non-specific binding up to 10-fold ^6^, which facilitates measurements that require activity of multiprotein complexes ^7,8^. However, this treatment may not be ideal for some applications as short DNA molecules and lipophilic dyes might bind to and diffuse through the detergent surface ^7^. In cases where very low concentrations of nucleic acids or proteins are used and tolerance to non-specific binding to surface is minimal, the sample chamber has been treated with a high concentration of blocking DNA or RNA (such as 1 mg/ml tRNA from yeast) ^9^.

Cleaning of the slides before passivation is another important step of surface preparation, especially when slides (such as quartz) are recycled by etching the previously treated surface to create a fresh layer with minimal fluorescence background. Sonicating the slides in potassium hydroxide (KOH) and acetone within an ultrasonic bath followed by piranha etching (a mix of sulfuric acid-H_2_SO_4_ and hydrogen peroxide-H_2_O_2_) are typical cleaning steps ^1,2^. As this summary illustrates, extensive efforts are required for cleaning and passivation of the surfaces before reliable measurements can be performed. One way to increase the yield of the slides is to segment them into multiple channels (used interchangeably with ‘chambers’), typically separated by double-sided tape. To enable buffer exchange through a channel, it is often necessary to drill (mechanically or using laser engraving) inlet and outlet holes, e.g. 6-12 holes for 3-6 channels, which adds to the labor and cost of each slide further motivating their recycling.

In cases where one or multiple parameters need to be varied systematically or a large number of conditions are to be tested, it is highly desirable to perform these studies under identical surface conditions for maximum consistency. Despite the time and effort spent to prepare the slides, variations in surface quality cannot be eliminated. Therefore, there are significant practical and fundamental advantages, especially in terms of reproducibility, in being able to use the same channel for as many measurements as possible. While this can sometimes be achieved by a judicious plan of experiments, such as starting from a low salt or protein concentration and gradually increasing the concentration in successive measurements, it is often necessary to reset the channel before new surface-attached molecules can be introduced. This would require breaking the bonds that immobilize the molecules of interest to the surface.

Despite variations among different surface passivation methods, biotin-streptavidin (or its analogs avidin or neutravidin) is the most frequently used surface attachment ^10^. This is typically achieved by mixing a few percent biotinylated PEG or BSA molecules with their unbiotinylated counterparts. Tetrameric streptavidin (with four biotin binding sites) is then introduced to the sample chamber and binds to the biotin-PEG or biotin-BSA molecules, which creates surface anchor points for biotinylated molecules (such as DNA, protein, lipid, or nanoparticles). Controlling the density of surface immobilized molecules is achieved by modulating the fraction of biotinylated PEG or BSA molecules and the concentration of streptavidin. While the high affinity biotin-streptavidin binding (dissociation constant K_d_ <10^-14^ M) makes it ideal for surface immobilization, the stability of the bond presents a challenge for resetting the sample chamber ^11^.

Several methods have been proposed to reset and recycle the sample chambers ^12^. For brevity, we will consider the particular case of DNA-protein interactions in which DNA molecules (or other types of nucleic acids or chimeras) are immobilized on the surface while protein molecules are in the solution. If single stranded DNA (ssDNA) is immobilized on the surface, one of the ends is typically modified with a biotin. In the case of immobilizing double stranded DNA (dsDNA) or partial duplex DNA (pdDNA), it is adequate to modify one of the strands with biotin. For these systems, a partial reset can be achieved by introducing 0.1% sodium dodecyl sulfate to the chamber, which removes the DNA-bound proteins while leaving the surface-bound DNA molecules intact ^12^. Another method to partially reset the system is to introduce a buffer with a high pH (such as 50 mM NaOH or LiOH) which breaks up the bonds between the two DNA strands and enables flushing out the non-biotinylated strand and any protein that might be bound to it ^12,13^. In the case of pdDNA constructs, the biotin is typically attached to the short strand while the longer strand contains an overhang that includes the protein interaction site. Therefore, removing the long, non-biotinylated strand also removes any bound proteins. Variations of these concepts and a combination of SDS and NaOH/LiOH treatment can be utilized to remove proteins and the non-biotinylated strand from a large group of nucleic acid constructs with different geometries (including pdDNA, R-loops, forked DNA, looped DNA, DNA with a bubble, kink, or various modifications). The common theme of these approaches is that one of the DNA strands (the biotinylated one) is left in the chamber and the second strand is introduced and annealed to the remaining strand at ambient temperatures. While this approach is effective in some cases, it has its shortcomings if the dsDNA, pdDNA or the protein-DNA complexes need to be treated in a particular manner before they are introduced to the sample chamber. For example, there will be challenges in applying this method if a protein-DNA complex needs to be pre-formed (before it is introduced to the sample chamber) by incubating the relevant molecules at elevated temperatures (to which the sample chamber cannot be subjected to), or the DNA constructs contain secondary structures that prevent them from effective hybridization at ambient temperatures. In such cases, having the capability to remove the entire DNA (including the biotinylated strand) or DNA-protein complex would be advantageous. While this would provide a fresh surface, it does require breaking the biotin-streptavidin bond which can be achieved by exposing the surface to a very high concentration basic solution (such as 7 M NaOH) ^12^. This approach was demonstrated to result in effective removal of neutravidin molecules and enabled recycling of a PEG coated sample chamber 10 times (by reintroducing neutravidin at every cycle) with similar levels of protein activity. However, it is not clear if the quality of surface passivation is impacted by exposure to such a strong reagent, and additional levels of surface passivation were suggested after this treatment.

In this study, we propose an alternative method to break the biotin-streptavidin attachment by using photocleavable biotin (PC-biotin) and ∼5-minutes of UV exposure ^14,15^. We demonstrate that this approach enables effective resetting and recycling of PEG coated surfaces for at least 10 cycles, without subjecting the surface to harsh chemicals or undesired ions. We tested different UV light sources (varying wavelength and power) and characterized the rate of bond breakage for different types of photocleavable biotin constructs (Schematic 1), which are all commercially available. We did not observe any degradation in surface quality during these recycling steps, as measured by non-specific binding of fluorescently labeled DNA and protein molecules. We also demonstrate the capability to recover DNA secondary structure formation and protein activity during these recycling rounds. We used a G-quadruplex (GQ) secondary structure and Replication Protein A (RPA) as the model systems in these studies ^16,17^. GQ structures are non-canonical secondary structures that are stabilized by Hoogsteen hydrogen bonds and stacking of planar tetrads ^18,19^. In addition to other environmental factors (including surface interactions), monovalent cations stabilize the GQ structures and impact their folding conformation in varying ways ^20– 23^, making them sensitive to the ions that are introduced to the sample chamber. RPA is the most abundant ssDNA binding protein in eukaryotes and is involved in a multitude of biological pathways ^24–26^. RPA is also known to destabilize GQ structures and stretch the unfolded ssDNA ^27^. We demonstrate effective recovery of both GQ folding and RPA-mediated GQ unfolding activity for at least 10 recycling rounds, further confirming the maintenance of surface quality.

## 2. Experimental Section

### 2.1 Photocleavable Biotin

As illustrated in Schematics 1, several versions of photocleavable biotin are available from commercial vendors, including Integrated DNA Technologies (IDT, Coralville, Iowa). PC-biotin is synthesized by inserting a photocleavable spacer between the biotin group and the DNA base. While the 5’ PC biotin is available as a standard modification, 3′ PC biotin, 5′ PC biotin Azide and 5′ PC biotin NHS can be attained as special order from IDT (which adds ∼$100 to the cost per strand). The molecular weight of biotin is 393.4 Da while that of 5’ PC biotin is 597.6 Da. The recommended UV wavelength for optimal cleavage is in the 300-350 nm range.

The UV cleavage of all these variants of PC-biotin required similar levels of exposure. However, we observed a significant reduction in the binding affinity of DNA oligos that contained PC-biotin compared those with standard biotin (called ‘biotin’ henceforth). While incubating 40 pM of an oligo with the biotin modification for 1 minute in a microfluidic chamber was adequate to obtain the desired coverage on the surface (∼ 300 molecules within a 100 μm x 50 μm imaging area), we needed to incubate 400-1000 pM of oligos modified with PC-biotin for several minutes to attain a similar coverage. We tested whether this difference is due to potential degradation during annealing or ambient light exposure during sample preparation (a total of four test groups as presented in Supporting Figure S1). We do not observe any variation in terms of affinity to bind to surface-immobilized streptavidin molecules among the four test groups, suggesting the oligos or the photocleavable attachment between biotin and the oligo do not degrade during annealing or sample preparation. Lower binding affinity of PC-biotin compared to biotin or degradation of oligos at an upstream step (before they are frozen for long term storage) are potential reasons we cannot rule out at this point. Nevertheless, the need to use higher concentrations of PC-biotin tagged oligos did not make a practical difference in terms of oligo consumption since the required concentrations (<1 nM) are at least two orders of magnitude lower than what is optimal for the annealing process (∼ 0.1-1.0 μM to increase the likelihood of hybridization). Typically, only a fraction of annealed constructs is used in any given experiment.

**Schematics 1.**
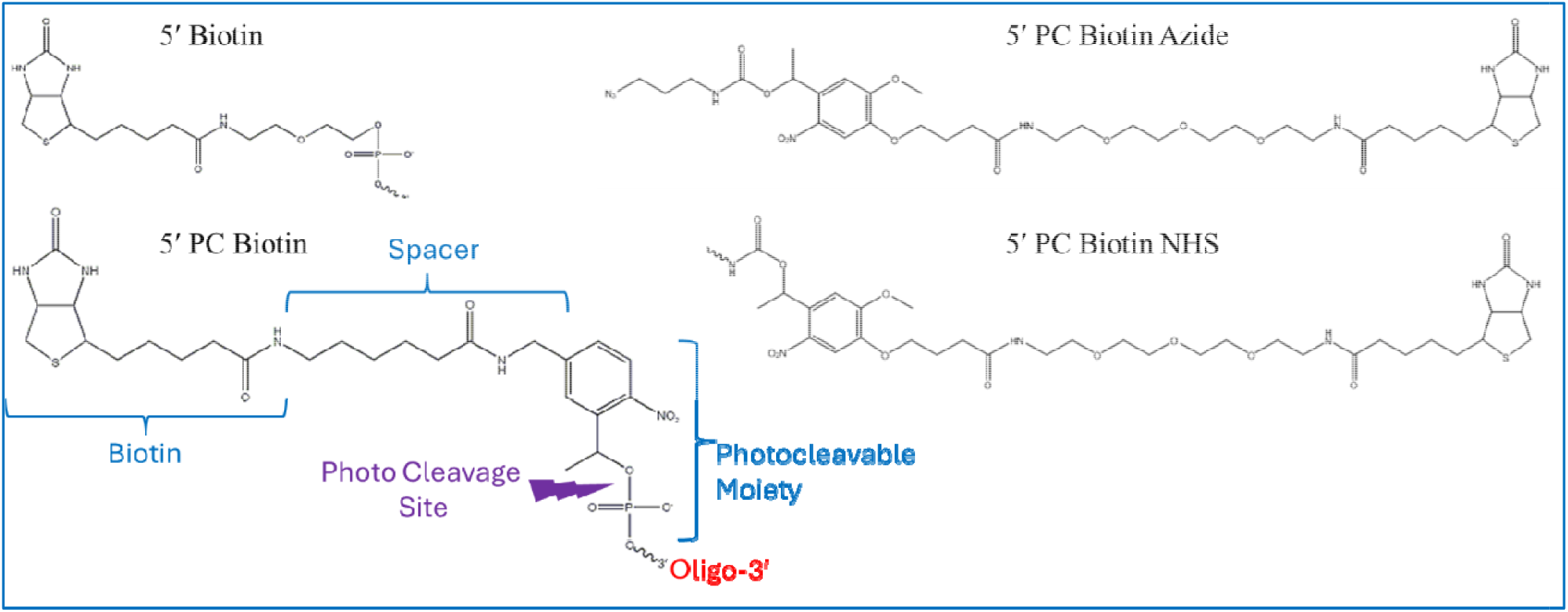
Chemical structures of biotin and photocleavable biotin modifications. The photocleavable moiety, photocleavage site and oligo attachment sites are indicated in the schematics for 5’ PC Biotin. The sequence of DNA constructs given Supporting Table S1 also lists which of these modifications were incorporated into each strand with cross-reference to relevant figure.

### 2.2 DNA and Protein Constructs

Schematics of the single molecule fluorescence and single molecule Förster resonance energy transfer (smFRET) assays, including the ssDNA and pdDNA constructs are shown in relevant figures. The sequences of all nucleic acids are given in Table S1. It is common practice to use pdDNA constructs in which the short strand (typically 18-30 nt long), which contains the biotin and one of the fluorophores (typically the acceptor), is kept constant while the long strand is modified according to the needs of the experiment. This enables using the same short strand in numerous experiments, e.g., we used the same short strand in 25 different constructs in a recent study ^28^. Therefore, attaching a photocleavable biotin to such short strands (instead of the biotin that is typically used) provides the capability of recycling the chamber and performing multiple measurements on different constructs under practically identical surface conditions. Human replication protein A (RPA) was purchased from Enzymax, LLC (Catalog # 61) and Cy3-labeled streptavidin (Cy3-streptavidin) from Acro Biosystems (Catalog # STN-NC114).

### 2.3 Single molecule fluorescence and smFRET measurements

A prism-type total internal reflection fluorescence (TIRF) setup built around an Olympus IX-71 microscope was used for imaging studies ^13^. The number of molecules that remain on the surface after a short period of UV exposure (Figure 1) were quantified by a peak finding algorithm based on imaging 35 different regions within the chamber (15-frame long movies with an exposure time of 100 ms/frame). The error bars in each data point represent the standard error obtained from these 35 measurements. Therefore, each region was exposed to laser beam excitation for ∼2 s, resulting in negligible photobleaching. A similar procedure was followed for smFRET measurements. The FRET distributions were determined by acquiring 30 short (15 frames) and 5 long (1500 frames) movies at different parts of the chamber before and after introducing RPA to the chamber. This was followed by exposing the slide to UV light for 5 minutes which removes all molecules from the surface. A fresh DNA sample was introduced to the sample and the procedure was repeated.

**Figure 1:**
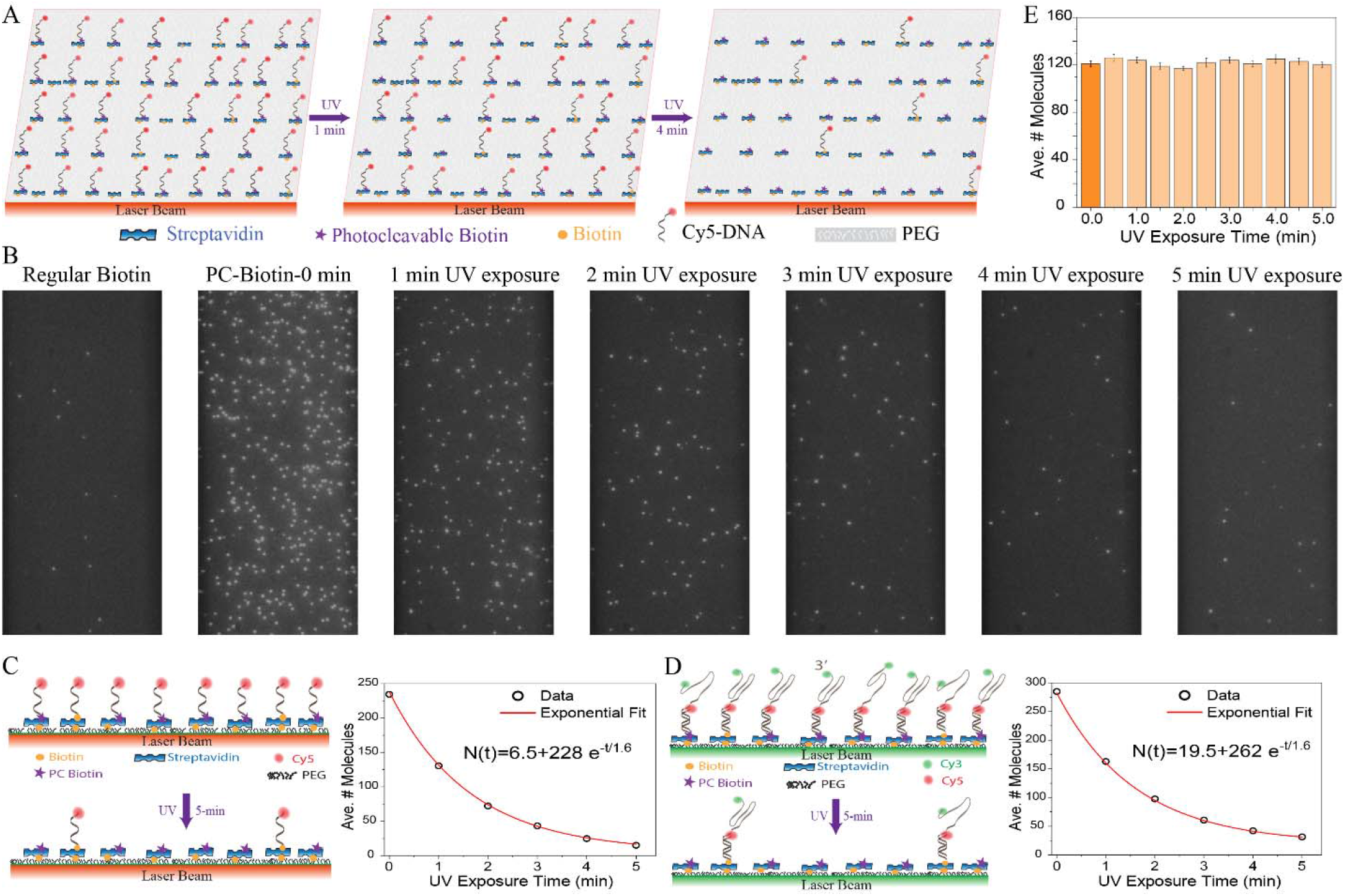
Characteristic time and efficiency of PC-biotin photocleavage. (A) Schematic of the photocleavage assay. Successive UV exposures result in gradual decrease in the number of surface-bound fluorescent molecules (Cy5-DNA tagged with PC-biotin) due to UV cleavage of the bond between PC-biotin and streptavidin. A small fraction of DNA molecules tagged with biotin (yellow circles) were mixed with those tagged with PC-biotin (purple circles) to serve as an identifiable background when all PC-biotin molecules are photocleaved after a total of 5-min UV exposure (right panel). (B) Representative images showing the decrease in the number of surface-bound molecules after UV exposure. The first image on the left shows the molecules attached via biotin. (C) Left: Schematics of the assay. Right: Number of surface-bound Cy5-DNA molecules as a function of UV exposure time shows an exponential decay with a characteristics time of 1.6 min. Five minutes is adequate to remove all surface-bound molecules. The flat background level around ∼10 molecules is due to the fluorescent molecules that are attached to the surface via biotin (those shown in the left-most image in (B)). (D) Left: Schematics of the assay. Right: Similar exponential behavior is observed for partial-duplex DNA molecules attached to the surface via a strand that is tagged with PC-biotin. The flat background level (around ∼20 molecules in this case) is due to the fluorescent molecules that are attached to the surface via biotin. (E) Similar experiments performed on molecules attached to the surface via biotin do not show a decrease in the number of surface bound molecules after multiple rounds of UV exposure (Supporting Table S2), demonstrating the absence of detectable UV-induced photobleaching.

To identify the surface and achieve proper focus, we added a small fraction (5-10%) of fluorescently labeled DNA molecules that were labeled with biotin, which does not photocleave. This is particularly important in cases where UV exposure removes all fluorescently-tagged molecules, introducing challenges and uncertainties for imaging the surface without reliable markers. Before adding the DNA molecules that contain PC-biotin in Figure 1 (or the molecules used to test non-specific binding in Figures 2-3), we quantified the number of the molecules attached via biotin. This number was then subtracted from the total number of molecules after molecules with PC-biotin (or those used for testing non-specific binding) were introduced.

**Figure 2:**
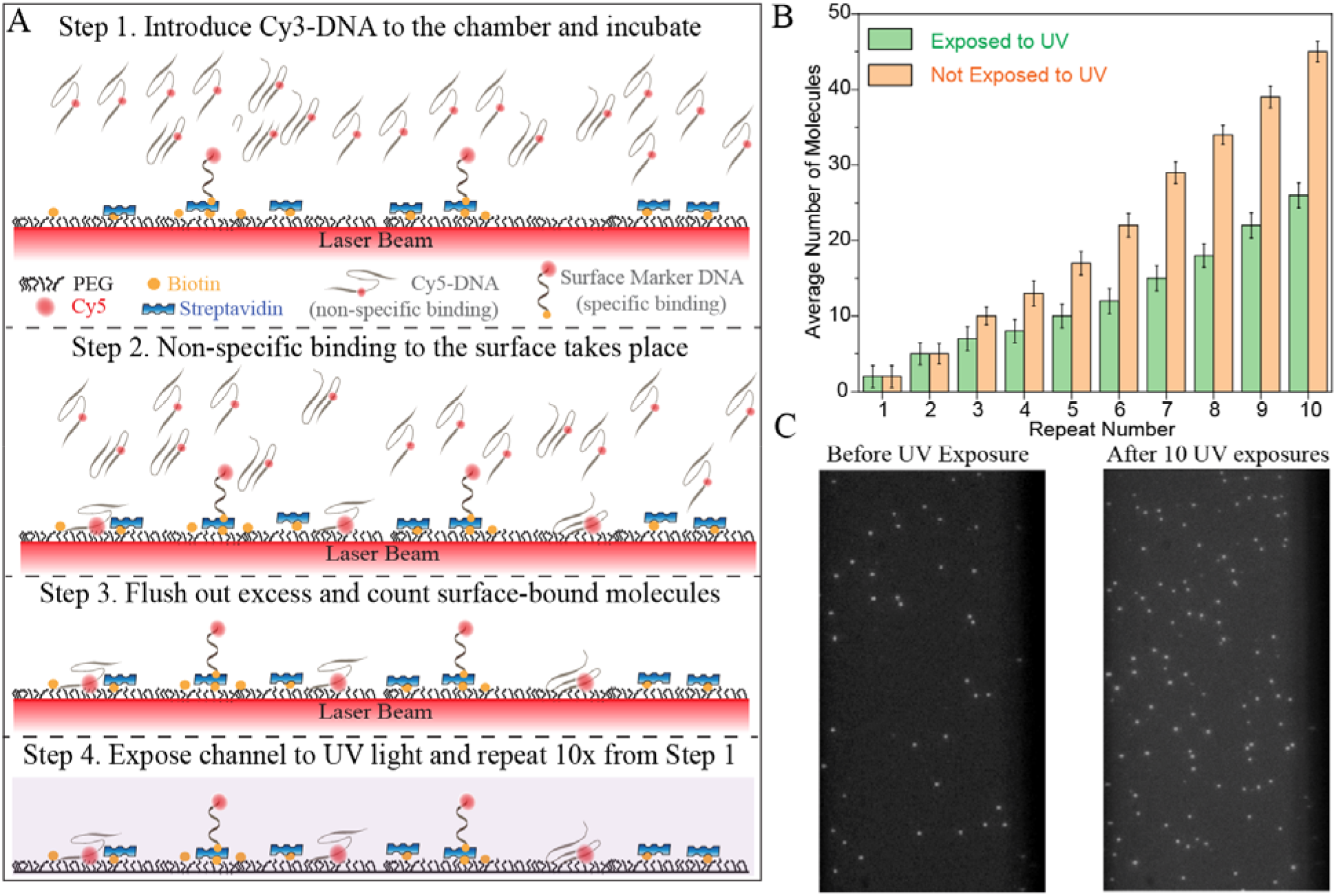
Impact of successive UV exposures on non-specific binding of Cy5-DNA. (A) Schematic of the assay where non-specific binding of Cy5-labeled DNA to a PEG surface is probed over 10 successive rounds of incubations. (B) Data on non-specific binding test performed on a UV-exposed or protected (not exposed to UV) chamber. The chambers were either subjected to 10 successive 5-min UV exposures or protected. Each data point represents the average number of molecules based on imaging 35 different regions on the surface. The error bars are standard error of these 35 measurements. As expected, the number of surface-bound molecules systematically increases with each round of incubation as non-specific binding is cumulative; however, the UV-exposed surfaces perform as well or better than the protected surfaces in each of these tests. One way ANOVA analyses within each group show the change in number of molecules with each repeat is significant (Supporting Table S3, p=0.001). T-test analysis comparing the UV-exposed and protected groups does not show significant difference between the two groups for Repeats 1-3 (p>0.05), while the differences are significant for Repeats 4-10 (Supporting Table S4, p<0.001). (C) Images showing the surface before UV exposure (just molecules that are specifically attached via biotin-streptavidin) and after 10 exposures (both specifically and non-specifically bound molecules are present). The increase in the number of molecules in the right image is due to cumulative non-specific binding over 10 rounds.

**Figure 3:**
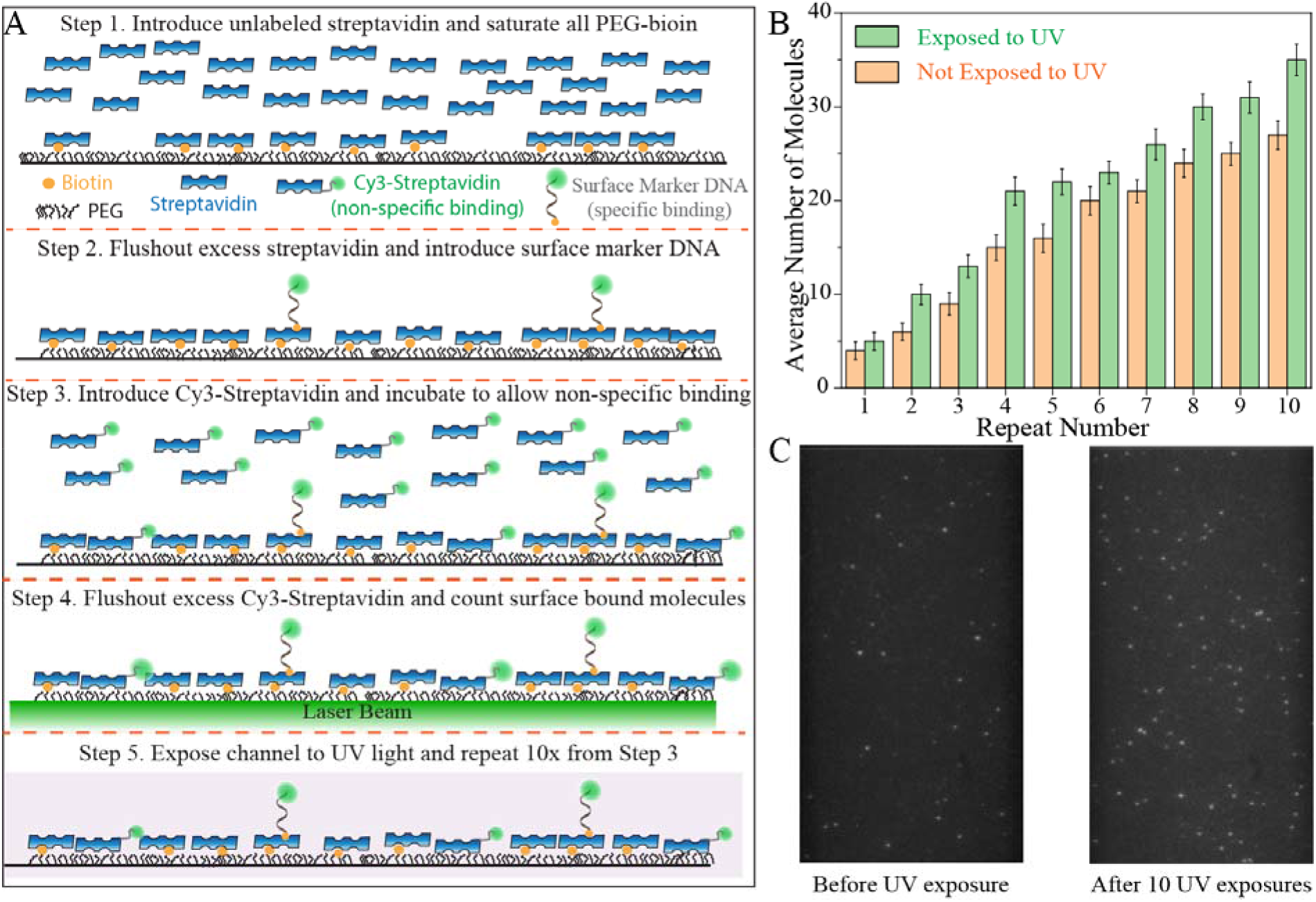
Impact of successive UV exposures on non-specific binding of labeled protein. (A) Schematic of the assay. Non-specific binding of Cy3-streptavidin to a PEG surface that is either subjected to 10 successive 5-min UV exposures or protected (not exposed to UV). The biotin-PEG molecules on the surface were saturated with unlabeled streptavidin before introducing Cy3-streptavidin by incubating a high concentration (100-fold excess) of unlabeled streptavidin. Therefore, only molecules due to non-specific binding attach to the surface. (B) Comparative data showing non-specific binding in a protected (not exposed to UV) or UV-exposed chamber. Each data point represents the average number of molecules based on imaging 30 different regions on the surface. The error bars are standard deviation of these 30 measurements. The number of surface-bound molecules systematically increases at successive cycles due to non-specific binding being cumulative. The channel exposed to UV and the protected channel are essentially indistinguishable in terms of non-specific protein binding, suggesting no detectable degradation in surface quality due to UV-exposure. One way ANOVA analyses within each group show the change in number of molecules with each repeat is significant (Supporting Table S5, p=0.001). T-test analysis comparing the UV-exposed and protected groups does not show significant difference between the two groups for Repeats 1 and 6 (p>0.1), while the differences are significant for the other repeats (Supporting Table S6, p<0.05). (C) Images showing the surface before UV exposure (just molecules that are specifically attached via biotin-streptavidin) and after 10 cycles of Cy3-streptavidin incubation and UV exposures (both specifically and non-specifically bound molecules are present). The increase in the number of molecules in the right image is due to cumulative non-specific binding over 10 rounds.

In order to characterize the time required to remove surface-bound molecules, we utilized fluorescently labeled ssDNA and pdDNA constructs that were modified with PC-biotin in either the 3′ or 5′ end. Using TIRF microscopy and appropriate laser beam excitation, we imaged different regions of the surface and counted the number of surface-bound molecules before and after exposing the slide to UV light for 30 s. We repeated this procedure until the number of molecules reaches background levels.

### 2.4 Sample Chamber Preparation

Laser drilled quartz slides and glass coverslips were thoroughly cleaned with acetone, 1M potassium hydroxide (KOH), and 20 min of piranha etching, followed by amino silane functionalization for 20-30 min. The slides and coverslips were then passivated with a mixture of mPEG (PEG-5000, Laysan Bio, Inc.) and biotin PEG (biotin-PEG-5000, Laysan Bio, Inc.) in 40:1 molar ratio. The PEG mixture was sandwiched between the slide and coverslip, which results in PEGylation of inner surfaces. These surfaces form the upper and lower surfaces of the microfluidic chamber; however, only the surface of the quartz slide is imaged in prism-type TIRF arrangement. After cleaning with distilled and deionized water and drying with nitrogen gas, the slides and coverslips were stored at -20 °C. Before the experiments, the slides and coverslips were thawed and passivated with 333 Da PEG for 30-45 min to increase the density of the PEG brush. The microfluidic chambers were created between the slide and coverslip with double-sided tape, followed by sealing the chamber with epoxy. Finally, the microfluidic chamber was treated with 2% (v/v) Tween-20 to reduce non-specific binding. After 10 min incubation, the excess detergent was removed from the chamber. For experiments where we tested non-specific binding (Figures 2-3), we further passivated the surface by incubating 4 μM transfer RNA (tRNA from baker yeast) for 10 minutes, followed by removing the excess molecules. After each UV-exposure step, we performed a buffer exchange to remove the free molecules (e.g., those dissociated from surface by UV-exposure) with a buffer that contains 50 mM Tris (pH 7.5), 150 mM KCl, and 2 mM MgCl_2_. To ensure all free molecules are removed, the buffer exchange was performed with at least 10-fold greater volume of buffer (>200 μl) compared to the volume of chamber (<20 μl).

### 2.5 UV Exposure

We tested several UV light sources, including commercially available UV lamps, flashlights, and a UV oven, to identify a reliable option. These light sources had different wavelengths (including 254 nm, 302 nm, 365 nm) and different powers. The option that worked best for single molecule applications was a UV lamp with a wavelength of 302 nm, power of 6 Watts and intensity of 1940 μW/cm^2^ (UVP, UVM-57 which is currently available through Analytik Jena Part Number: 95-0104-01). All data presented in the manuscript were obtained using this light source. While other light sources occasionally yielded effective cleavage of PC-biotin, the results showed significant variation between different trials and samples. We observed effective and rapid (∼30 sec) dissociation of PC-biotin when incubated within a UV Curing Oven (λ=365 nm, maximum intensity 250 mW/cm^2^, Edmund Optics Stock #26-822); however, the significant heat generated within the oven impacted the quartz slide and the double-sided tape that separated the channels from each other. Due to the lower cost, lower power, and reliable results obtained with the λ=302 nm UV lamp, we implemented this source for daily use. Supporting Figure S2 shows images of the sample chamber and UV lamp.

## 3. Results and Discussion

### 3.1 Determining the Cleavage Efficiency and Characteristic Time to Cleave PC-biotin

Fluorescently labeled DNA molecules were immobilized on the surface via PC-biotin and streptavidin attachment, in addition to a small fraction of molecules attached via biotin-streptavidin in order to serve as surface markers after the photocleavable molecules are removed (Figure 1A). The average number of molecules that remain on the surface after successive 30 s UV exposures (λ=302 nm, 1940 μW/cm^2^) was determined by imaging 35 different regions on the surface. Figure 1B shows images taken after systematically increasing UV exposure times, illustrating the reduction in the number of surface-immobilized molecules, which is well-described by an exponential decay with a characteristic decay time of 1.6 min (Figure 1C). For all samples we studied, 5 minutes of UV exposure was adequate to reduce the number of surface-bound molecules to background levels. Similar decay characteristics were observed for pdDNA constructs in which one of the strands was modified with PC-biotin (Figure 1D), suggesting the presence of a complementary strand does not inhibit the UV cleavage. To check whether the UV exposure photobleaches the fluorescent molecules, hence impacts the observed decay in the number of surface-bound molecules, we repeated the same measurements with DNA molecules that were attached to the surface via biotin-streptavidin, which should not cleave when exposed to UV light. Figure 1E shows that the average number of such molecules does not change over 10 UV exposure cycles (see Supporting Table S2 for ANOVA analysis supporting the variations not being statistically significant), confirming that the decrease in the number of molecules attached via PC biotin is due to cleavage, rather than UV-induced photobleaching.

### 3.2 Impact of UV Exposure on Non-Specific Binding of DNA and Protein Molecules

We investigated how the PEG surface quality is impacted by successive 5-min UV exposures by quantifying non-specific binding of fluorescently labeled DNA and protein molecules. If surface quality degraded due to successive UV exposures, we expect to observe an increase in non-specific binding. To quantify non-specific binding, we used fluorescently labeled DNA or protein molecules. These molecules were not biotinylated, hence could not specifically bind to the surface.

Figure 2 shows results of 10 successive measurements probing non-specific binding of Cy5-DNA molecules (10 nM concentration) to the surface. After quantifying the number of molecules in each measurement, the slide was exposed to UV-light for 5-minutes and a fresh batch of Cy5-DNA molecules was added to the chamber. As a control, we repeated the same procedure in different channels that were not exposed to UV light. As non-specific binding is cumulative, we expect an increase in the number of surface bound molecules; however, significant degradation in the surface quality due to UV exposure should result in higher levels of non-specific binding. The UV-exposed channels performed as well or in some cases better than the control channels (a surprising result that requires for further investigation), suggesting 10 successive rounds of UV exposure (a total of 50 minutes) do not degrade the surface. Supporting Table S3-S4 provide T-test and one-way ANOVA analysis, respectively, on these data. Figure 2C shows representative images of the surface before UV exposure and after 10 rounds of introducing DNA molecules and exposure to UV. The bright spots in the left image are the molecules that are introduced to serve as surface markers, attached via biotin-streptavidin as described in Experimental Section. These molecules serve as a constant background and their number is subtracted from the total number of molecules, resulting in the non-specifically bound molecules presented in Figure 2B. We also performed similar measurements with 1 nM DNA, which is more representative of the DNA concentration used to immobilize molecules on the surface and did not observe detectable non-specific binding at this relevant concentration even after 10 cycles of UV exposure (Supporting Figure S3).

Figure 3 shows measurements in which we tested the impact of UV exposure on surface quality as gauged by non-specific binding of Cy3-streptavidin. In order to prevent specific binding of Cy3-streptavidin to the PEG-biotin molecules, we pre-incubated the sample chambers with saturating levels of non-fluorescent streptavidin (concentration∼10^-7^ M compared to K_D_∼10^-14^ M). After all available biotin molecules were blocked and excess streptavidin was removed, we introduced biotinylated and Cy3-labeled DNA (Cy3-DNA) molecules to serve as surface markers and quantified their number. As saturating levels of streptavidin were attached to the surface, the number of these markers were higher than the typical numbers in other measurements (Figure 3C, left image). We then introduced 1 nM Cy3-streptavidin to two sample chambers and incubated them for 5 minutes. After removing the excess Cy3-streptavidin by purging the chamber, we determined the number of surface bound molecules in each chamber. Subtracting the number of specifically bound Cy3-DNA molecules from the total number of molecules results in the number of non-specifically bound Cy3-streptavidin molecules. We then exposed one of the sample chambers to UV light for 5 minutes while the other chamber was protected from exposure. We repeated this procedure of introducing 1 nM Cy3-streptavidin and UV exposure/protection 10 times so one of the chambers was exposed to 50 minutes of UV light while the other was protected. Figure 3 shows these data where a systematic increase in the number of surface bound Cy3-streptavidin molecules is observed in both the UV-exposed and the protected chamber. As non-specific binding events are cumulative, this trend is not surprising. However, the number of non-specific Cy3-streptavidin events is similar in UV-exposed and protected channels and the variation between the two cases is within the differences observed among difference channels (see Supporting Table S5-6 for relevant statistical analyses). Even after 10 rounds of Cy3-strepatividin incubation and UV exposure, the number of non-specifically bound molecules remains modest. To illustrate, in a typical measurement ∼300 surface bound molecules per imaging area is targeted while ∼35 non-specifically bound molecules were accumulated on the surface after 10 rounds of incubation.

### 3.3 Recovery of DNA secondary structure and protein activity

Having established that no detectable degradation is observed in the PEG surface due to UV-exposure, we then tested whether biological activity can be recovered in chambers that have been recycled by previous UV exposure. In the experiments presented in Figure 4, we investigated recovering of G-quadruplex folding pattern and of RPA-mediated GQ unfolding activity. The pdDNA constructs used in these studies contain an ssDNA overhang that folds into GQ, which compacts the overhang. Given the positions of the donor and acceptor fluorophores, the compaction of the overhang results in a higher FRET state. On the other hand, unfolding of the GQ and stretching of the ssDNA overhang by RPA results in a lower FRET state. The smFRET studies presented in Figure 4 illustrate the reproducibility of this process, i.e., folding of the GQ and its destabilization by RPA, over 10 cycles in which all surface-bound molecules are removed, and new DNA are introduced, followed by introducing (Fig. 4A). The panels on the left in Fig. 4B show a single high FRET peak representing folding of the GQ construct (before introducing RPA) and the panels on the right show a new low FRET peak representing RPA-bound state after RPA is incubated in the chamber. As illustrated with these data, both the smFRET histograms representing the GQ-only state and those representing the protein activity are highly reproducible even after 10 rounds of recycling.

**Figure 4:**
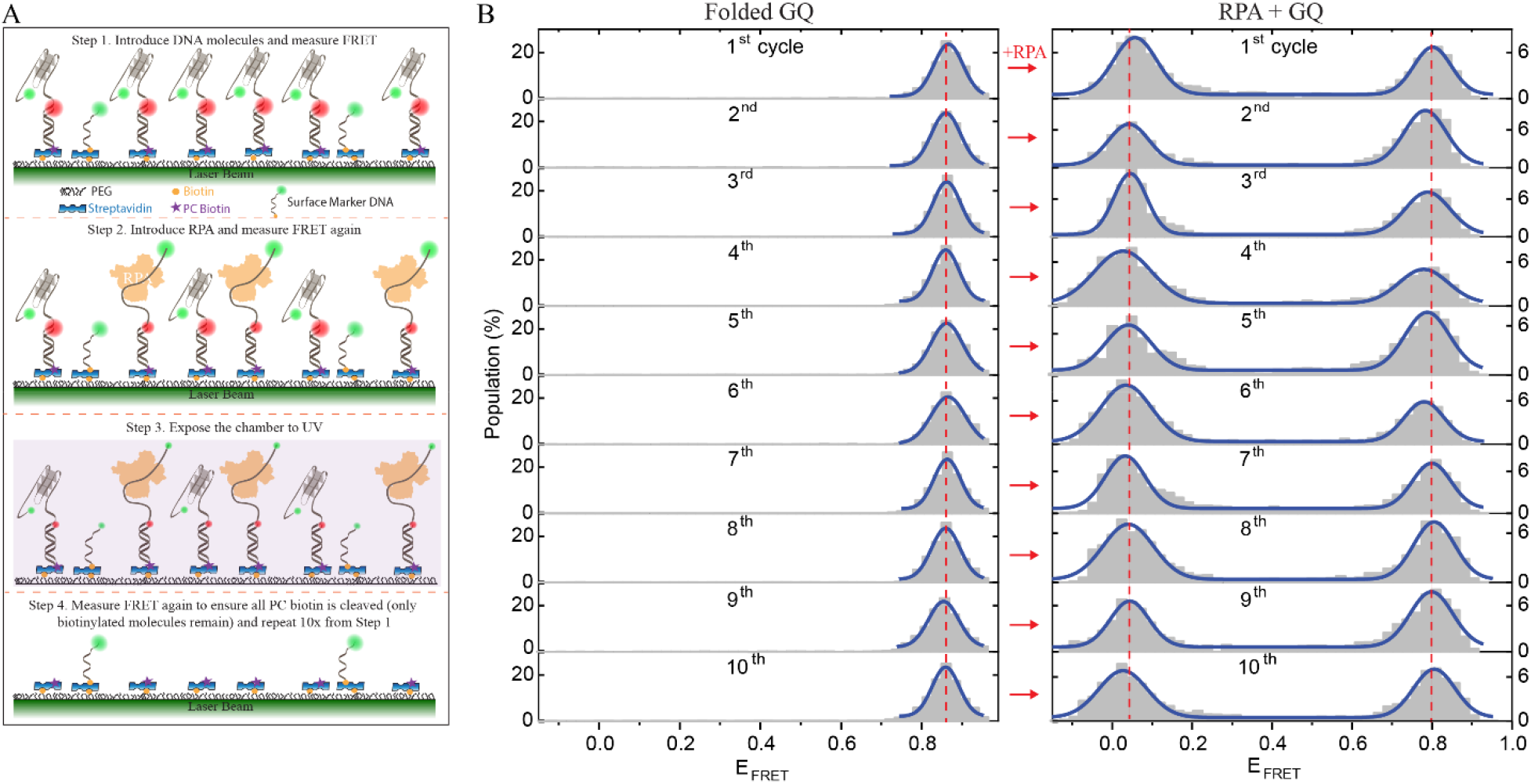
Reproducibility of G-quadruplex folding and RPA activity. **(A)** Schematics of the assay. Partial duplex DNA constructs with an overhang that can form a G-quadruplex structure are immobilized on the surface using PC biotin. The compaction of the overhang by GQ results in a high FRET peak. Binding to and unfolding of the GQ by RPA results in an extended overhang, hence a lower FRET state. After GQ folding and RPA mediated unfolding are recorded, the channel is reset by subjecting it to 5-min UV exposure. A fresh batch of pdDNA is then introduced and imaged, followed by introduction of RPA and imaging of the molecules. This cycle is repeated 10 times. A small fraction of biotinylated Cy3-labeled ssDNA molecules are kept in the channel to identify the surface and ensure all molecules attached with PC-biotin are removed. **(B)** Results of GQ folding (left panel) and RPA-mediated unfolding (right panel) at 10 successive rounds of recycling. The similarity of the folding pattern and level of protein activity suggests the recycling process enables consistent and reproducible recovery of the folding pattern and protein activity for at least 10 recycling processes. The solid lines are Gaussian fits to the FRET distributions.

## 4. Conclusions

We demonstrate that the attachment between photocleavable biotin and streptavidin can be practically broken, which enables complete removal of these molecules from the surface. A 5-minute exposure to light from a standard UV lamp (with a specific wavelength) was adequate to remove all molecules attached to the surface with PC-biotin. It is then possible to introduce new DNA molecules and repeat new measurements within the same chamber, a process that can be repeated at least 10 times without detectable degradation in the surface. The practicality of the approach, common use and affordability of UV lamps, and the short duration of the required exposure (which might be shortened with higher power UV sources) are some of the attractive aspects of the approach. Unlike alternative methods of surface regeneration, UV exposure does not require introducing strong reagents or potentially undesirable ions to the chamber. While an initial investment is required to switch from strands with biotin to those with PC-biotin, the time and resources saved by recycling the sample chamber and the consistency enabled by performing many experiments within the same chamber quickly offset this initial investment.

## Supporting Information

The Supporting Information is available free of charge at ….

Sequences of DNA constructs, report of statistical analysis, additional data, and image of the UV exposure setup

## Author Contributions

J.A. and H.B. conceived the concept. J.A. and T.A. conducted the single molecule experiments. J.A. and S.-G.K. analyzed the data and M.L.K. purified and labeled some of the DNA constructs. J.A., S.-G.K. and H.B. prepared the figures. H.B. wrote the manuscript and all authors reviewed and approved the manuscript.

## Acknowledgements

NIH [1R15GM146180 to H.B.] funded this work. J.A. is funded by the Deanship of Scientific Research at Northern Border University, Arar, KSA through the project number NBU-SAFIR-2024. Funding for open access charge: NIH Grant and Kent State University Research Council.

## Notes

The authors declare no competing financial interest.

## Table of Contents Graphics

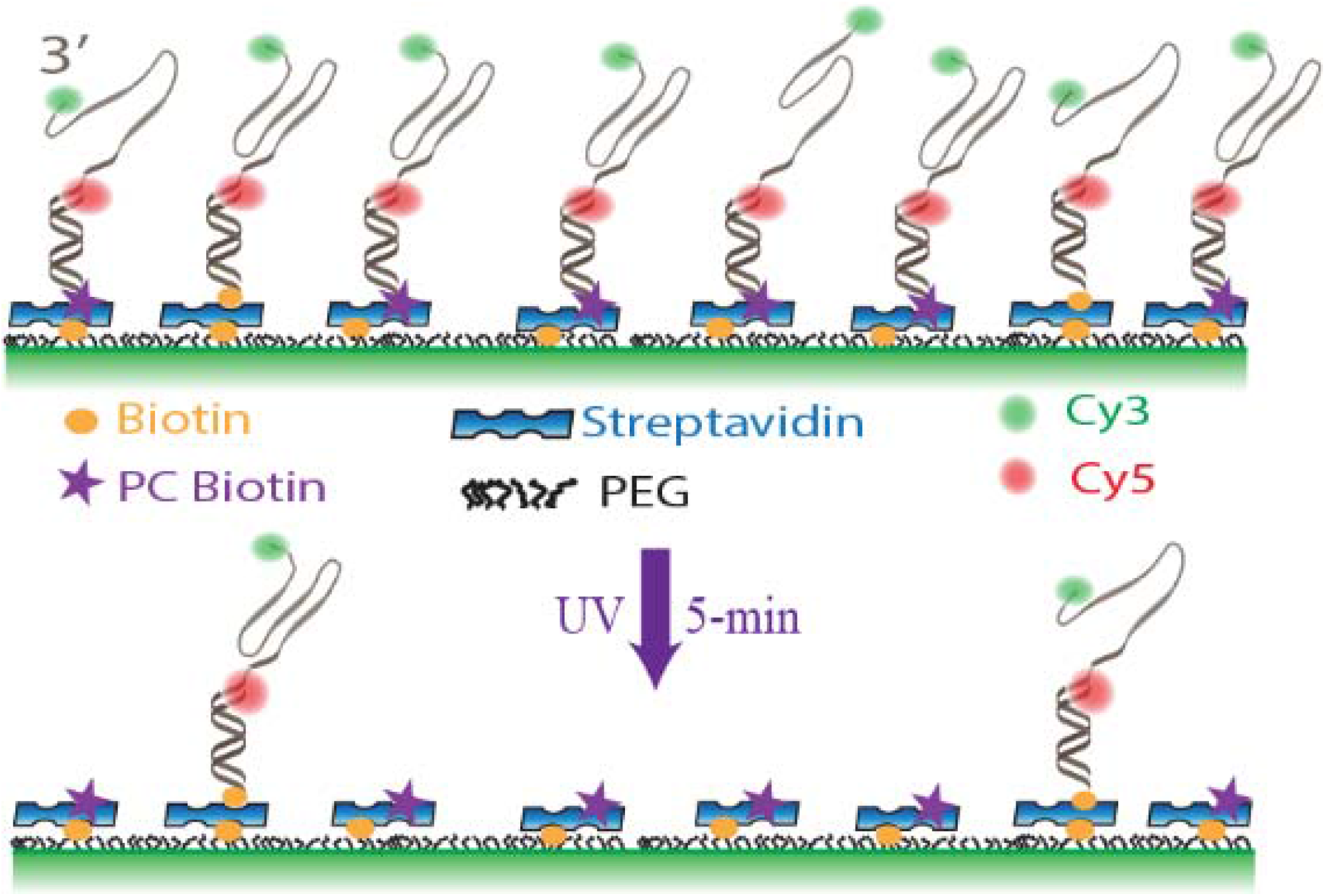

