## Supplementary Information for "Reusable Microfluidic Chambers for Single-Molecule Microscopy"

**Table S1**: Sequences of DNA strands used in this study.

| Strands | Sequence in 5′–3′ | Used in | Complement/Notes |
|---|---|---|---|
| Stem-30-Biotin-Cy5 | /Cy5/CCC TAA CCC TAA GCC TCG CTG CCG TCG CCA-Biotin | Figs.1C, 1D, 2 | DNA Marker in Fig. 2 |
| Stem-30-PCBioN- Cy5 | /Cy5/CCC TAA CCC TAA GCC TCG CTG CCG TCG CCA-PCBioN | Figs. 1C, 1D | Removed by UV exposure in Figs. 1C-D |
| Stem-30-Biotin-Cy3 | /Cy3/CCC TAA CCC TAA GCC TCG CTG CCG TCG CCA-Biotin | Fig. 3 | DNA Marker in Fig. 3 |
| Stem-18-Cy5 | GCC TCG CTG CCG TCG CCA /Cy5/ | Fig. 2 | Used to quantify non-specific binding in Fig. 2 |
| Stem-18-PCBioN-Cy5 | PCBioN-GCC TCG CTG CCG TCG CCA/Cy5/ | Fig. 4 | Short strand of GQ forming pdDNA |
| 2-Layer-3nt loop-Cy3 | /Cy3/TTG GTT TGG TTT GGT TTG GTT TGG CGA CGG CAG CGA GGC | Fig. 4 | Long strand of GQ forming pdDNA. Complements Stem-18-PCBioN-Cy5 |
| 3GTract-Cy3 | TGG CGA CGG CAG CGA GGC TTA GGG TTA GGG TTA GGG TTA G/Cy3/ | Fig. 1D | Long strand of pdDNA in Fig. 1D. Complements Stem-30-PCBioN- Cy5 and Stem-30-Biotin-Cy5 |

**Statistical Analysis**
In Tables S2-S5, we presented T-test and Analysis of Variance (ANOVA) tests performed for the data presented in Figures 1-3.

**ANOVA Analysis Table for Figure 1**
**Table S2:** One-way ANOVA test statistics comparing the average number of molecules within each group in Fig. 1E.

| Parameter | ANOVA test | Comment |
|---|---|---|
| UV exposure time | $F(10, 264) = 0.978$, $p=0.462$ | No Significant difference |

**ANOVA Analysis Table for Figure 2**
**Table S3:** One-way ANOVA test statistics comparing the average number of molecules within each group (Exposed to UV and not exposed to UV) in Fig 2B.

| Condition | ANOVA test | Comment |
|---|---|---|
| Exposed to UV | $F(9, 340) = 48.238$, $p=0.001$ | Significantly different |
| Not Exposed to UV | $F(9, 340) = 218.075$, $p=0.001$ | Significantly different |

**T-test Analysis Table for Figure 2**
**Table S4:** T-test analysis illustrating the statistical significance of the differences in the average number of molecules between two groups (exposed to UV vs not exposed to UV) in each repeat in Figure 2B.

| Repeat number | T-Test | Comment |
|---|---|---|
| 1 | $t(34) = -0.428$, $p=0.671$ | No Significant difference |
| 2 | $t(34) = -0.109$, $p=0.913$ | No Significant difference |
| 3 | $t(34) = -1.894$, $p=0.066$ | No Significant difference |
| 4 | $t(34) = -3.090$, $p=0.003$ | Significantly different |
| 5 | $t(34) = -4.497$, $p<0.001$ | Significantly different |
| 6 | $t(34) = -6.054$, $p<0.001$ | Significantly different |
| 7 | $t(34) = -10.161$, $p<0.001$ | Significantly different |
| 8 | $t(34) = -10.764$, $p<0.001$ | Significantly different |
| 9 | $t(34) = -11.041$, $p<0.001$ | Significantly different |
| 10 | $t(34) = -11.497$, $p<0.001$ | Significantly different |

**ANOVA Analysis Table for Figure 3**

**Table S5:** One-way ANOVA Test Statistics comparing the average number of molecules within each group (Exposed to UV and not exposed to UV) in Figure 3B.

| Condition | ANOVA test | Comment |
|---|---|---|
| Exposed to UV | F(9, 340)= 57.145, p=0.001 | Significantly different |
| Not Exposed to UV | F(9, 340)= 44.894, p=0.001 | Significantly different |

**T-test Analysis Table for Figure 3**

**Table S6:** T-test analysis illustrating the statistical significance of the differences in the average number of molecules between the two groups (exposed to UV vs not exposed to UV) in each repeat in Figure 3B.

| Repeat number | T-Test | Comment |
|---|---|---|
| 1 | t(34)= -0.418, p=0.678 | No Significant difference |
| 2 | t(34)= -4.283, p<0.001 | Significantly different |
| 3 | t(34)= -2.559, p=0.015 | Significantly different |
| 4 | t(34)= -2.754, p=0.009 | Significantly different |
| 5 | t(34)= -2.824, p=0.007 | Significantly different |
| 6 | t(34)= -1.515, p=0.138 | No Significant difference |
| 7 | t(34)= -2.336, p=0.025 | Significantly different |
| 8 | t(34)= -2.722, p=0.010 | Significantly different |
| 9 | t(34)= -2.850, p=0.007 | Significantly different |
| 10 | t(34)= -3.661, p<0.001 | Significantly different |

**Impact of Annealing and Light Exposure on Surface Coverage of PC-Biotin Tagged Oligo**

As stated in the Experimental Section, at least an order of magnitude higher concentration of DNA constructs tagged with PC-biotin is needed to attain similar surface coverage compared to those tagged with biotin. This difference is potentially due to degradation of the PC-biotin attachment to the DNA. To test whether annealing of the oligos or exposing them to ambient light during sample preparation causes such degradation, we performed controlled studies with four groups of DNA constructs prepared in different ways:

Group 1. Annealed and exposed to ambient light
Group 2. Annealed but protected from ambient light (Dark)
Group 3. Not-annealed and exposed to ambient light
Group 4. Not-annealed but protected from ambient light (Dark)

Figure S1 shows average number of surface bound molecules (top column graphs for 200 pM and 400 pM DNA) and example surface images when samples were prepared under these conditions. For each condition, we imaged 30 different regions on the surface and quantified the average number of surface bound molecules. As illustrated, we do not observe a significant difference between these different sample preparation conditions and conclude that annealing of the samples and exposing them to ambient light during experiments are not the cause of the difference between oligos tagged with PC-biotin or biotin. We speculate reduced affinity of PC-biotin (compared to biotin) to streptavidin as the cause for the observed difference; however, we can not also rule out potential degradation in an upstream step of the process before they are frozen for long term storage (e.g., shipping at ambient temperature). The vendor (IDT DNA) has not at this point tested the relative binding affinities of biotin and PC-biotin (private communication with IDT-DNA).

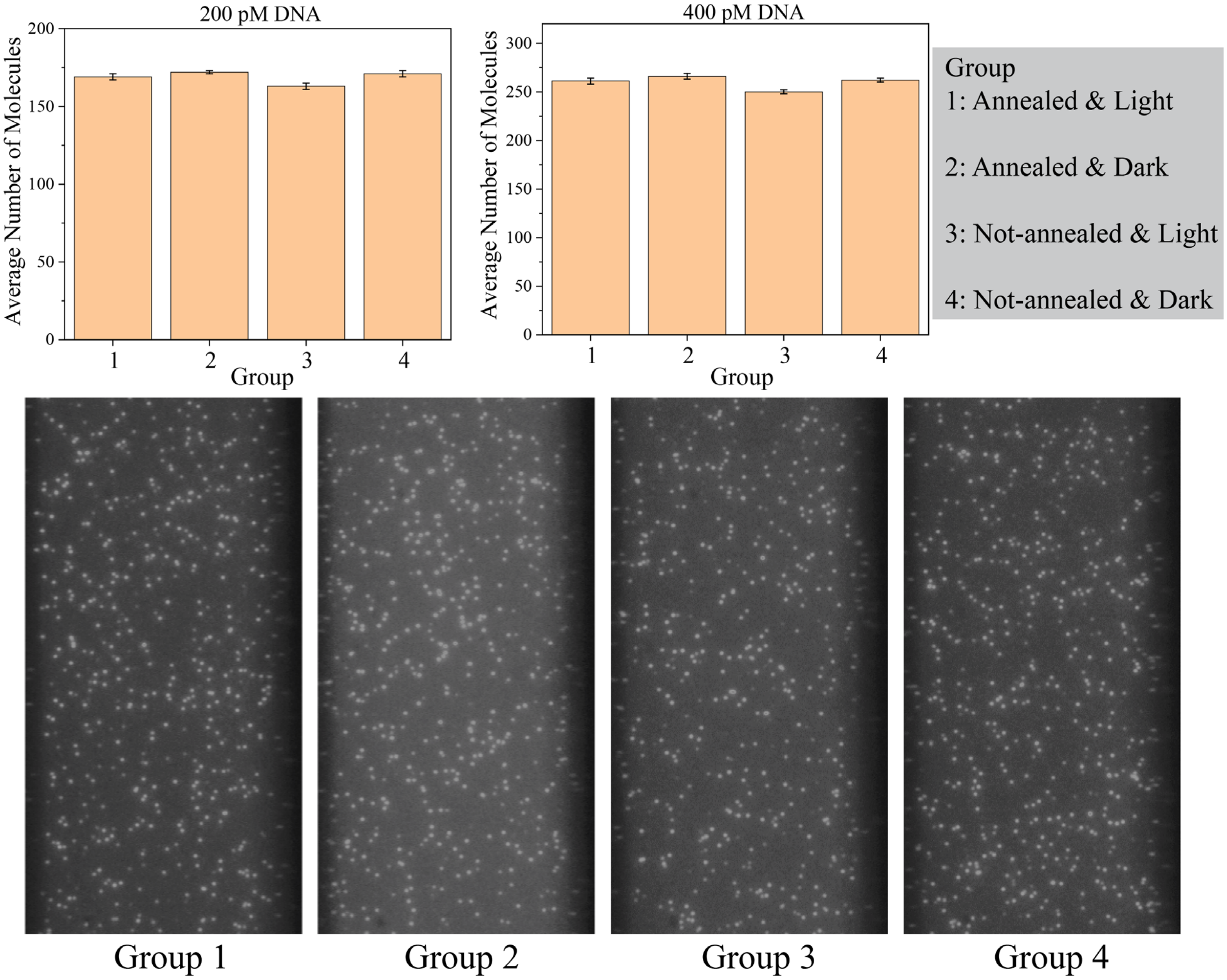


**Figure S1:** Surface coverage and average number of streptavidin-bound molecules tagged with PC-biotin and prepared under four different conditions, two different DNA concentrations for each case. (Top) Column graphs for DNA constructs at 200 pM (left) or 400 pM (right) concentration. 30 different regions on the surface were imaged in each case. Each column represents the average number of molecules based on these 30 measurements and the error bars are standard error of these measurements. (Bottom) Example images showing the surface coverage for each group.

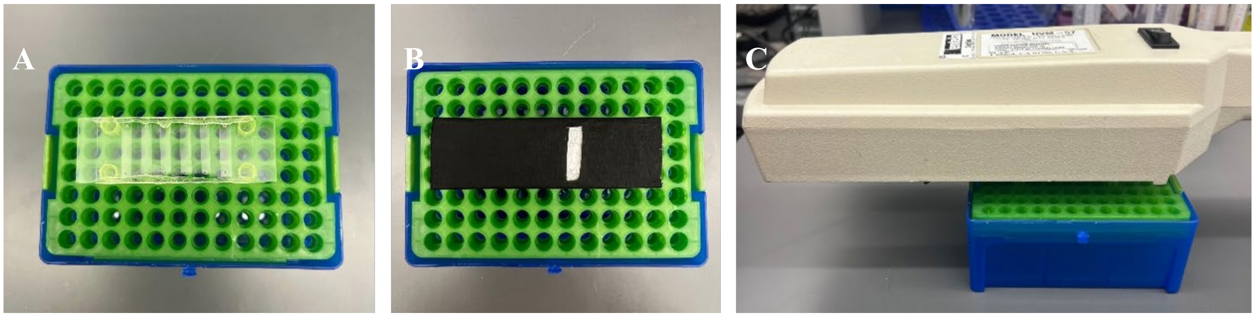


**Figure S2:** Images of sample chambers and UV lamp. (A) A close-up view of the slide with six sample chambers. (B) A picture showing an exposed sample chamber while others are protected with black cardboard. (C) A picture demonstrating the positioning of the UV-lamp directly above the slide with a gap of less than 1 cm between the beam and the slide.

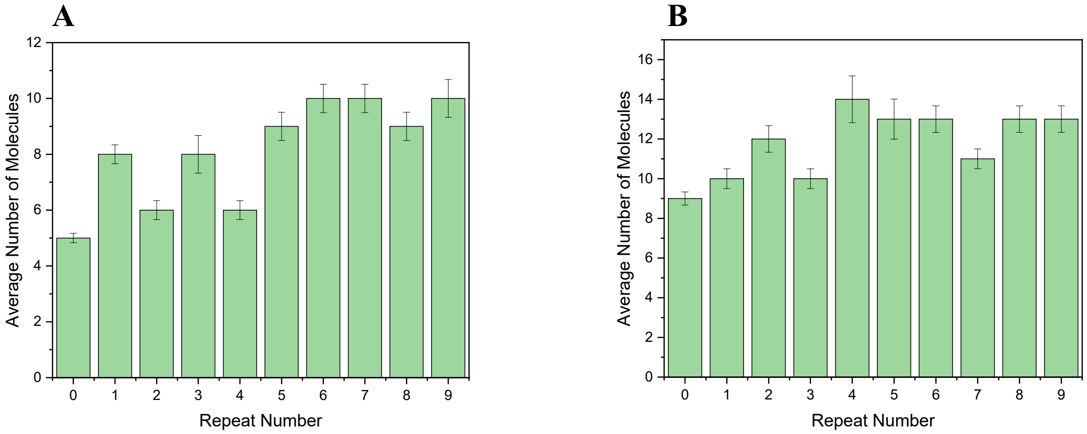


**Figure S3:** (A) and (B) Two independent tests of non-specific binding for 1 nM Cy5-labeled DNA to the PEG surface. Both chambers were exposed to UV light for nine cycles (Cycle 0 is before UV exposure). Each data point represents the average number of molecules based on imaging 35 different regions on the surface. The error bars are standard error of these 35 measurements. The number of specifically bound molecules (5 in **A** and 9 in **B**) is not subtracted from these data as the number of non-specifically bound molecules reduces to 0-5 range, which sometimes is smaller than the standard error for the respective cycle.
